# Functional requirements of intentional control over the integrated cortico-thalamo-cortical and basal ganglia systems using neural computations

**DOI:** 10.1101/2020.07.20.211425

**Authors:** Sébastien Naze, James Kozloski

## Abstract

Large scale brain models encompassing cortico-cortical, thalamo-cortical and basal ganglia processing are fundamental to understand the brain as an integrated system in healthy and disease conditions but are complex to analyze and interpret. Neuronal processes are typically segmented by region and modality in order to explain an experimental observation at a given scale, but integrative frameworks linking scales and modalities are scarce. Here, we present a set of functional requirements used to evaluate the recently developed large-scale brain model against a learning task involving coordinated learning between cortical and sub-cortical systems. The original Information Based Exchange Brain model (IBEx) is decomposed into functionally relevant subsystems, and each subsystem is analyzed and tuned independently and with regard to its relevant functional requirements. Intermediate conclusions are made for each subsystems according to the constraints imposed by these requirements. Subsystems are then re-introduced into the global framework. The relationship between the global framework and phenotypes associated with Huntington’s disease is then discussed and the framework considered in the context of other state-of-the-art integrative brain models.

## I. Introduction

**T**he Information-Based Exchange Model (IBEx [9]) is composed of multiple subsystems arranged both anatomically and functionally in a brain-inspired manner to simulate brain activity at multiple scales. The aim of the present work is to reproduce aspects of the experiment described in [8] using the IBEx model. The original experiment consists of an auditory tone being presented to a mouse having two electrodes implanted in the motor cortex, whereby the pitch of the tone depends on the filtered electrodes’ signals. The experiment is illustrated in **Figure 1**. In the successful realization of the task, the animal learns to control the pitch of the auditory tone by modulating its cortical neuronal firing, without initiating movement. The study shows that pharmacological blockade of cortico-striatal glutamatergic projections prevents the animal from learning the task.

**Figure 1:**
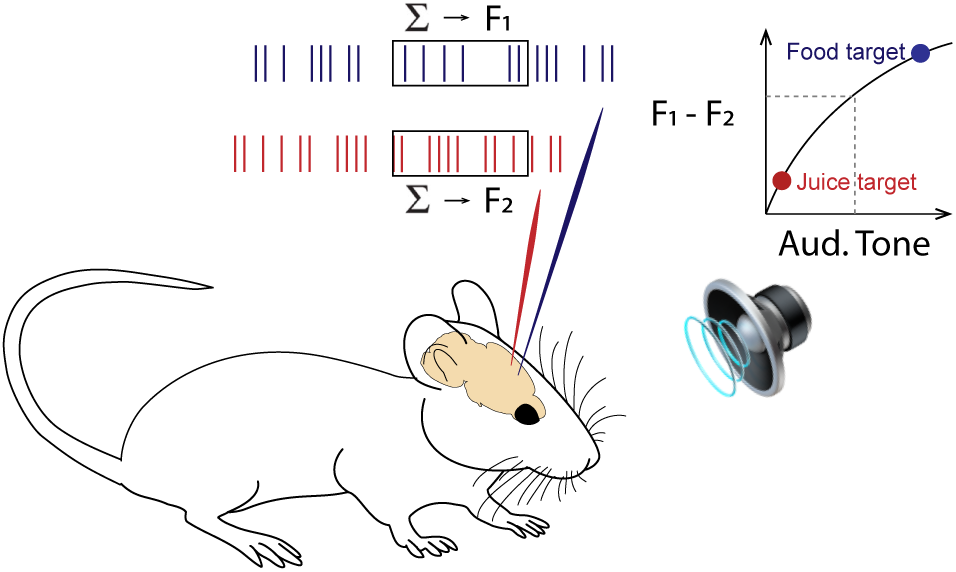
Overview of the Brain-Computer Interface (BCI) experiment to be reproduced in-silico. Two electrodes located in motor cortex M1 record neural activity in the form of multi-unit spiking activity. Firing rates F_1_ and F_2_ are computed over 1s windows and are used to control the pitch of an auditory tone, which is fed back to the animal. Target points are set for high and low pitch with associated food and juice rewards. In the reference study, the animal learns to control the tone by modulating its neural activity to receive the reward.

We present a set of functional requirements necessary to complete this task in the context of a theoretical model integrating cortico-thalamo-cortical and basal ganglia information processing [9]. Those requirements imply 1) the encoding of an auditory stimulus occurs in sensory cortices; 2) the stimulus is transmitted to other cortical areas; 3) the global cortical state is integrated in the striatum; and 4) the selection and sustenance of a given cortical state (i.e. a motor command) occurs via striato-pallido-thalamic projections onto frontal cortices.

In order to satisfy those requirements, we decompose the IBEx model into the following subsystems: 1) Stimulus encoding and decoding in superficial layers of cortex; 2) Stimulus propagation across cortical areas; 3)Striatal network properties allowing the integration of cortical state into an action plan; 4) Striato-pallido-thalamic gating for cortical states selection based on basal ganglia processing.

This manuscript focuses on either cortico-thalamo-cortical dynamics alone, or the striato-pallido-thalamic gating system projecting to the cortex, whereby striatal inputs are manually set for experimental purposes. As such, in contrast to the closed-loop system introduced in [9], our study uses an open-loop, since cortico-striatal projections and their integration are set aside for future work.

## II. Methods

The computational model has been described in detail in [9], and a diagram of the architecture is presented in **Figure 2**. In this section we provide a definition of the different components the system.

**Figure 2:**
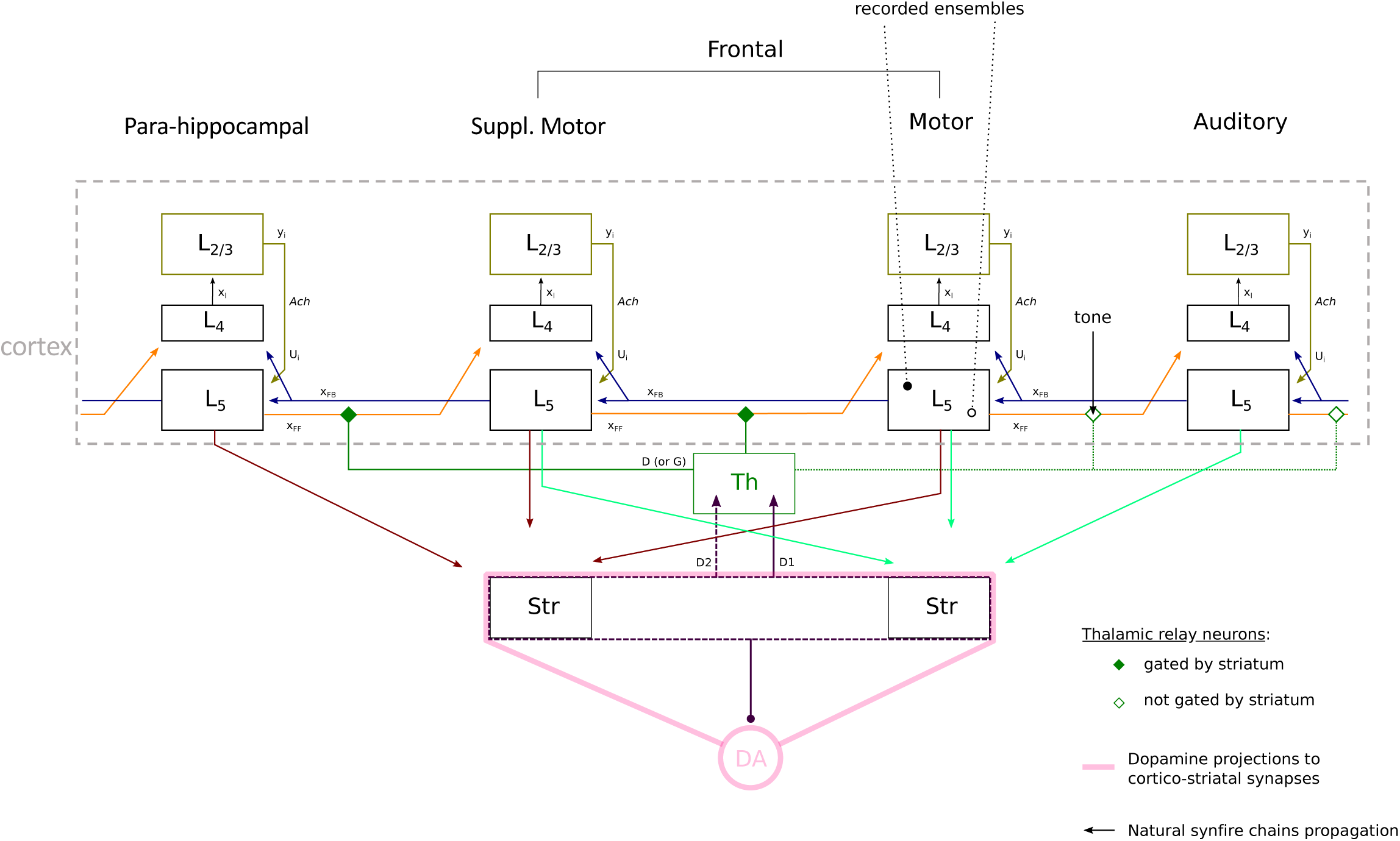
Diagram of the neuro-anatomically inspired system to reproduce the experiment in-silico. The model comprise 4 cortical areas (columns called Para-hippocampal, Suppl. Motor, Motor and Auditory), the thalamus (Th), striatum (Str), and dopamine neurons from the Substantia nigra pars compacta (DA). Each cortical area is composed of a supra-granular layer (L2/3, L4) modeled as an information maximization system, and an infra-granular layer (L5) modeled as a subnetwork of self-organized recurrent units. Key variables and parameters are shown for congruence with mathematical equations and reference code. X_FF_: feedforward cortico-thalamo-cortical activity; X_FB_: feedback cortico-cortical activity; Ach: cholinergic gain; D1/D2: direct/indirect basal ganglia pathway; D/G: thalamic gating of cortical activity.

### i. Model

The IBEx model comprises a cortico-cortical system producing synfire chains undergoing entropy maximization, a thalamo-cortical system gating feedforward information flow in cortical areas, a basal ganglia network of inhibitory neurons implementing the gate function, which is applied to thalamic relay neurons via striato-pallido-thalamic projections.

A detailed description of each subsystem of the model is as follows:

#### Information maximization algorithm (Infomax) for L2/3

The Infomax model [13, 14, 15, 10] is a three-staged network used in [9] to couple thalamic relay neurons to neocortex L4, L2/3 and L5, whereby L5 neural activity is mediated via cholinergically modulated drive from L2/3.

Stage one receives the rate vector 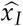 from an ensem-ble of time averaged thalamic spike trains and computes the zero-mean input vector 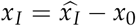, where 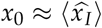 is learned at the learning rate 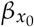

Stage two computes the sum of weighted inputs to stage three, *u*≡ *Cx*_*i*_, where *C* is a fully-connected weight matrix. In addition, each stage two unit computes an element of the output vector *y, y*_*i*_ = *σ*(*u*_*i*_) where *σ*(.) denotes the logistic transfer function 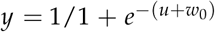 and *w*_0_ is an adaptive output bias learned by 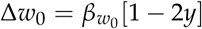.

Stage three computes an entropy maximizing term which is fed back to stage two to update *C* with learning rate *β*_*C*_:

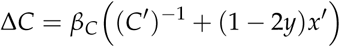

Since the inversion of *C* requires the knowledge of the full matrix (which is not biologically plausible) a local learning rule has been derived in [15] which uses an auxiliary vector *v* and lateral connections weights *Q* to approximate (*C*^*′*^)^−1^:

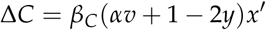

whereby *v* is computed by power iteration over 8 time steps,

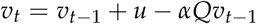

where *t* is a timer step and *α* is a positive number chosen to ensure that *αv* converges, such that 0 ≤ *α* ≤ 2/*Q*^+^, where *Q*^+^ is the largest eigenvalue of *Q* [14, 15].

The cholinergic modulation acts as a gating function over cortical layer 5, with gain *Ach* ∈ [0, 1] by

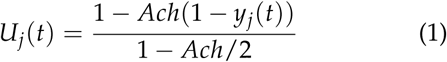

so that *U* ∈ [0, 2] for *y* ∈ [0, 1].

#### Self-Organized Recurrent Network (SORN) for L5

SORN [12] is a binary spiking neural network model made of excitatory and inhibitory units with multiple structural, synaptic, and homeostatic plasticity rules.

The network has been shown to autonomously and robustly generate synfire chains through a spontaneously generated feed-forward network composed of multiple sub-networks or clusters [27], while reproducing the approximately log-normal distribution of cortical synaptic strength observed experimentally [25]. Given the state of excitatory unit *x*(*t*) and inhibitory unit *y*(*t*) at time step *t*, the neuronal dynamic at the following time step reads:

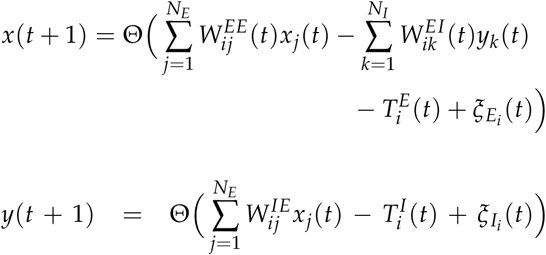

where Θ is the Heavyside step function, *N*_*E*_ and *N*_*I*_ the number of excitatory and inhibitory units, *T*^*E*^ and *T*^*I*^ the threshold values of excitatory and inhibitory units drawn from a uniform distribution in the interval 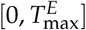 and 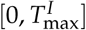. 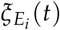 and 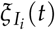 are gaussian noise process with mean *µ*_*ξ*_ = 0 and variance *σ*_*ξ*_^2^ ∈ [0.01, 0.05].

Spike-timing dependent plasticity (STDP) operates on excitatory-excitatory synapses by affecting the weight matrix *W*^*EE*^ through a window of one time step:

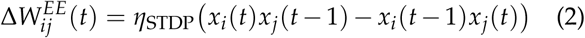

where *η*_STDP_ is the learning rate. Synaptic weights of negative values are eliminated by the rule, and to compensate for it, a new connection of weight 0.001 is created with probability *p*_*c*_ = 0.2 between two unconnected pairs. Efferent excitatory-excitatory synaptic weights are normalized by multiplicative scaling

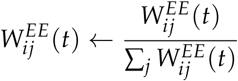

The target firing rate 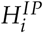 of an individual unit is maintained through a intrinsic plasticity rule acting over the threshold 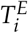:

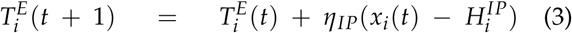

where *η*_*IP*_ is the adaptation rate and 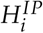 is drawn from a uniform distribution in [*µ*_*IP*_ − *σ*| *HIP, µ*_*IP*_ + *σ*|*HIP*].

Finally, inhibitory to excitatory synaptic weights evolve under an inhibitory STDP (iSTDP) rule

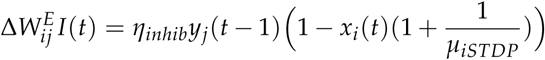

where *η*_*inhib*_ is the adaptation rate and *µ*_*iSTDP*_ = 0.1. Default parameters are *N*_*E*_ = 400, *N*_*I*_ = 100, *η*_*IP*_ = 0.01, 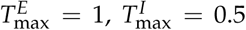, *µ*_*IP*_ = 0.1, *σ*_*HIP*_ = 0, *η*_*inhib*_ = 0.001, 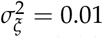, and *η*_*STDP*_ = 0.004.

After an initial period of about 1,000,000 time steps, the SORN system alone converges to a stable feedforward network, which is then used in the global IBEx architecture to implement the cortico-cortical system. In the IBEx system, all synaptic weights are still under the effect of STDP, but a connectivity mask *A*_0_ is introduced to maintain the nature of intra- and inter-area cortical connections constrained from cytoarchitectonics projections exposed in [9] (see **Figure 2**).

#### A recurrent inhibitory network of Fitzugh-Nagumo neurons for the striatum

Medium spiny neurons (MSNs) represent the large majority of neurons in the striatum (about 90% [22]). Our striatum dynamics focus on a network composed solely of this neuronal type, which we modeled using a Fitzhugh-Nagumo neuron model [4, 7]. A reduced version of the Hodgkin-Huxley model of the action potential [6] it comprises only two dimensions, which reproduce the fundamental properties of membrane excitability via state variables *v* and *w*, representing the membrane potential and a recovery constant:

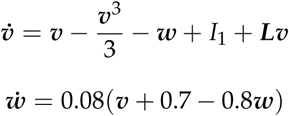

where *I*_1_ in a baseline input current, and ***L*** is a lateral inhibitory weight matrix. In the context of the IBEx model [9], ***v*** is re-interpreted as the bursting potential, whereby a single action potential is re-interpreted as a burst of action potentials.

We configured and tested 2 connectivity profiles for this inhibitory network:

- A randomized network with connection probability *p*_*s*_ = 0.35 implementing a *winnerless competition*, with bidirectional connection as in [9], and without as in [26].
- A distance-based connectivity network using a discretized kernel based on the gamma distribution as in [21] implementing a localized stable *winner-take-all* dynamic or unstable transitions between several localized activity bumps according to the nature of the input (Section iv).

#### Integrate-and-Fire (IAF) neurons with rebound-bursting properties for the Substantia nigra pars compacta (SNc)

The reinforcement learning system in the original IBEx model was implemented through a striato-nigral loop, whereby the Fitzhugh-Nagumo MSNs projected to Mihalas-Niebur IAF neurons [17]. Reinforcement learning occurs at cortico-striatal synapses and differentially on D1- and D2-receptor expressing neurons [9]. In the context of our study, this part of the system was discarded and we focused only on cortical or cortio-thalamo-cortical dynamics and the striato-thalamo-cortical network.

#### A rate model for thalamic integration and binary gating of L2/3 input based on striatal dynamics

The model of thalamic processing is composed of 3 parts: 1) the gating of frontal cortices based on the basal ganglia output via striato-pallidal projections; 2) the integration of primary sensory inputs to the feedforward cortical input stream; 3) the temporal integration of feed-forward and feedback cortical streams.

1. The striato-pallidal gating of thalamic relay neurons projecting to frontal cortices is referred as the forward driving gating function *G*(*v, x, t*) [9]. D1 neurons exerting a *go* action and D2 neurons a *no-go* action via *G*, which is computed using a pallido-thalamic adjacency matrix *D*, comprising 1 for direct pathway inputs, −1 for indirect pathway inputs, and 0 for all other inputs. The bursting of striatal MSNs are represented by a half-wave rectified function ℋ of the burst potential ***v***(*t*) from the Fitzhugh-Nagumo model:

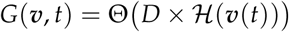

where Θ(·) is the Heaviside step function.
The feed-forward thalamo-cortical stream ***x***_*FF*_(*t*) is computed based on the summed spike train of SORN excitatory neuron activity ***x***(*t*) and a full-rank random mixing matrix *M*_*FF*_ required by infomax such that

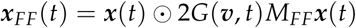

where ⊙ denotes element-wise multiplication.
2. A sensory input *s*(*t*) can be incorporated to the feed-forward stream (when applicable) by aggregating an input vector to the feed-forward vector *x*_*FF*_(*t*) = *x*_*FF*_(*t*) + *s*(*t*).
3. The temporal integration of feed-forward and feedback streams is performed through the sum of each stream ***x***_*TH*_ (*t*) = ***x***_*FF*_ + ***x***_*FB*_, followed by the spike train sum over a thalamic integration temporal window *τ*:

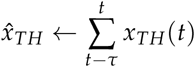

Note that some modifications from the original model were carried out in order to satisfy the functional requirements, mainly:

- The striatal network structure can now be set to either random (with or without bidirectional connections), or distance-dependent (using a nearest-neighbours connectivity kernel). The effect of such connectivity change is investigated in section iv.
- Striato-pallido-thalamic projections now follow a topographic organization [20], implemented as center-surround projection: D1-MSN disinhibitory projections onto the ventral thalamus (direct pathway) are spatially converging from pallidum to thalamus (i.e. center); while the D2-MSN inhibitory projections to the ventral thalamuc through SNc (indirect pathway) are spatially divergent (i.e. surround).
- Thalamic gating of the feedforward corticothalamo-cortical pathway affect granular and agranular cortices differently: as suggested in [2], thalamo-cortical projections are more diffuse in granular cortices compared to agranular cortices. In the revised model, thalamic inputs to agranular cortices project to all layers of cortex, while projections to granular cortices are L4 specific.

### ii. Auditory stimulus

The sensory stimulus *s*(*t*) is inspired by the auditory processing of the cochlea and auditory brainstem, in that we consider the vector of neural activity of the auditory cortex A1 to implement a tonotopic map of receptive fields, causing neurons to respond based on the pitch of an auditory tone [23]. For simplicity, we divide the auditory cortex neuronal units into 3 sets responding to high, medium and low frequency tones. The auditory tone is first integrated from the thalamo-cortical projections by element-wise multiplication of the auditory tone to the feed-forward pathway after thalamic filtering. A stimulus is presented every 2 to 5 seconds and lasts 1 to 2 seconds.

### iii. Post-processing

The original output of SORN takes the form of hundreds of spiking units, with connectivity that self-organize into feedforward subnetworks of different sizes. In the IBEx model, each subnetwork is interpreted as a cortical region [9]. Here, we derived a metric to quantify and visualize the amount of information carried by the signal in each cortical region by transforming SORN unit spike trains into firing rates. Each SORN unit spike train is time averaged using a sliding rectangular window of 100 ms, and a singular value decomposition (SVD) is applied to the series of all units’ firing rates of a given feedforward subsystem, reflecting the dynamics of infragranular layers (i.e. L5) of a cortical area. The 3 largest singular values of the overall activity are used to create a 3-dimensional space onto which the trajectory of the cortical activity can be projected to obtain a low-dimensional representation of the hundreds of spiking neurons’ signals. The process is illustrated in **Figure 3**.

**Figure 3:**
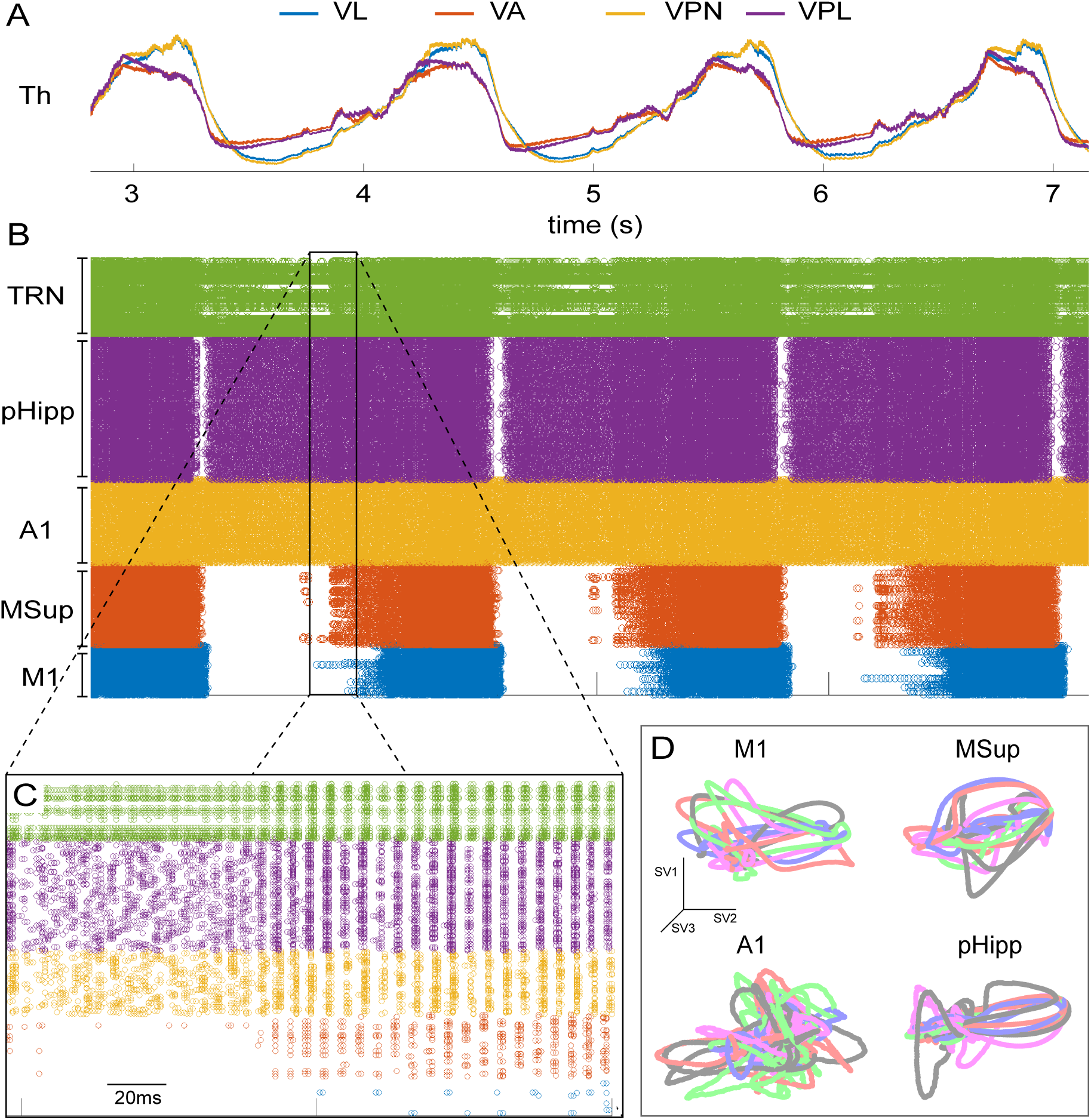
From SORN spiking units to cortical trajectories. A) Thalamic activity (Th) of 4 thalamic nucleus (VL: ventro-lateral; VA: ventro-anterior; VPN: ventro-posterior; VPL: ventro-posterior lateral) projecting feedforwardly to the 4 cortical areas of SORN (M1: primary motor; MSup: supplementary motor; A1: auditory; pHipp: para-hippocampal). B) SORN units spike train after self-organization into 4 clusters which are interpreted as cortical regions (legend for regions is consistent with A). C) Magnified view of SORN units’ spike trains. D) SVD outputs for each SORN subsystem, interpreted as cortical trajectories for each brain region. N.B. trajectories’ colorcode is different from A-B-C. Here for visualization purpose, each color represents a 2s segment of cortical activity.

### iv. Simulation platform

Simulations were performed using the IBM Model Graph Simulator [11] on a Lenovo P910 workstation (12 cores at 3.2GHz, 64GB RAM). A 1,000,000 time steps simulation equivalent of 1000s of simulation time took an average of 15 minutes to run. Prototyping of the model was performed

## III. Results

The IBEx model was decomposed in multiple subsystems to satisfy functional requirements of aspects of the learning task exposed in [8]. In the first part of this section, we present the baseline activity of the original model and a dimensionality reduction framework to analyze its dynamics, focusing on coupled and decoupled cortical systems. In the second part, we identify the functional requirements for each subsystems and tune the models parameters relevant to satisfying those requirements. Visualizations of the model’s outputs in the form of low-dimensional trajectories capturing the overall dynamics of the infra-granular cortical information flow is presented, under several architectural modifications.

### i. Baseline cortico-cortical processing

We started by isolating the cortical system from the rest of the model. There are 2 elements in the cortical subsystem. First, a Self-Organized Recurrent Network (SORN) which is initialized with 4 subnetworks forming synfire chains [27] in what is referred to as the Layer 5 (L5) cortico-cortical feedback pathway. It spans 4 cortical areas [9] whereby the direction of the information flow goes from agranular to granular cortices [24]. Second, for each cortical area, an infomax model [14] representing L2/3 receives L5 SORN input in the opposite direction than the synfire chain in what is described as the feed-forward cortical pathway [9], see **Figure 2** for a diagram of the system. In each cortical area, the output of infomax *U*_*i*_ modulates the SORN subnetwork activity by applying a multiplicative coupling to the SORN input *x*_*FB*_, scaled by the parameter *Ach* representing cholinergic modulation strength (Eq.1).

Figure 4 shows the low-dimensional trajectories of L5 activity (SORN) in M1 and values of the 10 first singular values (SVs) of a 10s simulation with and without infomax (L2/3) coupling through multiplicative modulation. We observe that without coupling (*Ach* = 0), SORN trajectories form a low-dimensional manifold in the 3-dimensional space of its largest singular values (see Methods section for details of the construction of the low-dimensional projection). The low information content of the SORN activity is further supported by the distribution of SVs, whereby the first SV (SV1) captures about 67% of the variance of the signal. In contrast, when the coupling between infomax and SORN is strong (*Ach* = 1), SV1 accounts to only 20% of the signal and the following SVs distribution is more uniform. This is also reflected in the low-dimensional trajectories, which span a larger space and do not follow a repetitive path.

Interestingly, a slow rhythm in the delta frequency range (1-4Hz) is observed in the thalamo-cortical system in both conditions, albeit more strongly present in the infomax decoupled system (*Ach* = 0) than coupled (*Ach* = 1). **Figure 3A** shows the average timeseries of thalamic averaging, with clear oscillatory activity in the delta band.

**Figure 4:**
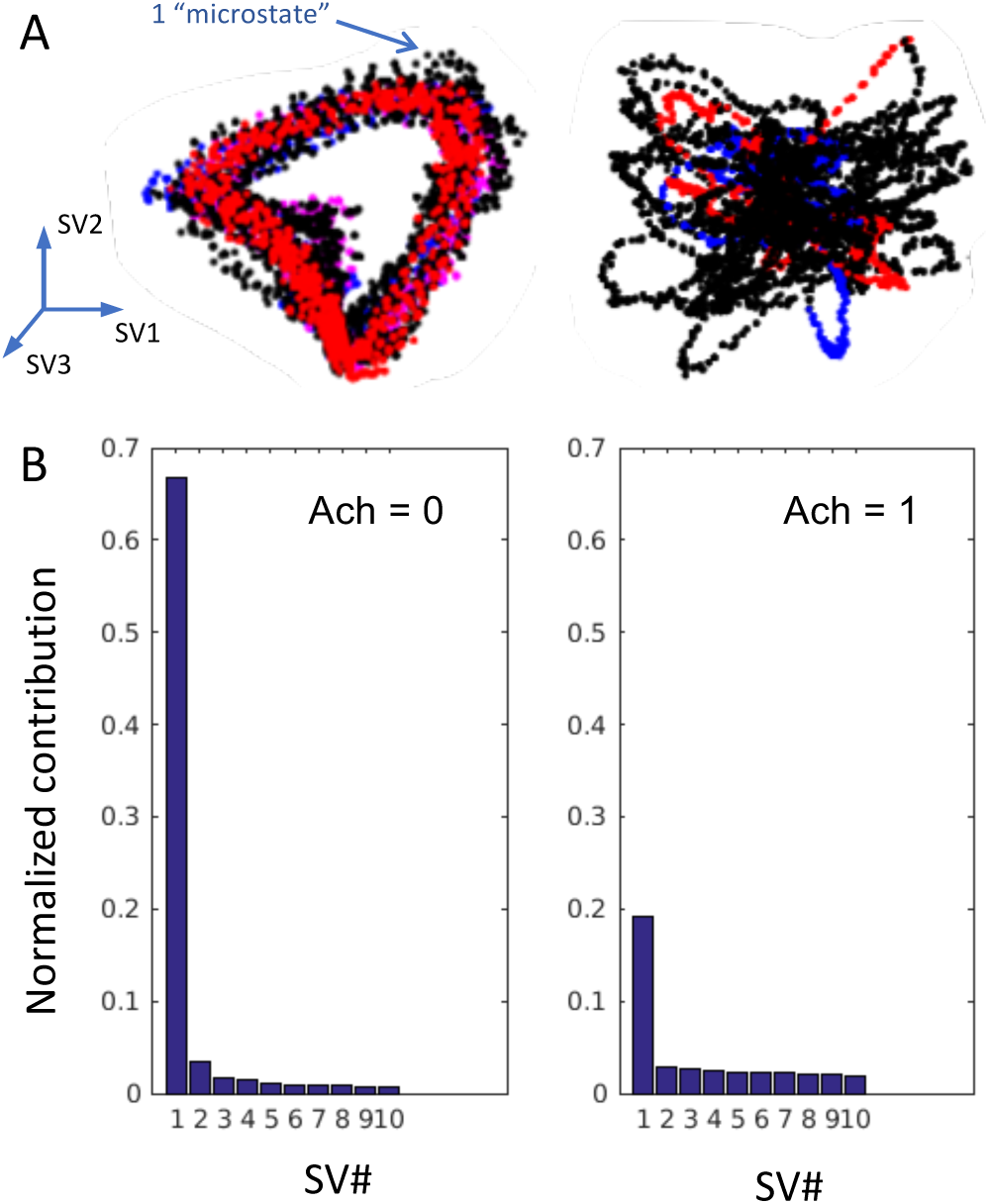
Effect of information maximization from L2/3 on L5 self-organized recurrent network SVD. Projection of modeled Auditory cortex A1 firing rate activity onto low dimensional space described by the 3 largest components of the singular value decomposition (SVD), without (left) and with (right) coupling between L2/3 to L5. Blue and red colors indicate are each a 2s second of activity in the low dimensional space of cortical activity. Distribution of the first 10 singular values (SV) of L5 firing rate activity for A1 region, L2 normalized to unity across all SVs (number of SVs vary by region).

### ii. Stimulus encoding in L2/3

When sensory inputs are introduced through the thalamo-cortical projections to L2/3 via L4, the modelled auditory stimulus is incorporated as an infomax input, whereby the information content of the L2/3 response about the input is maximized. The output is then used to modulate SORN activity as shown in Section i. The encoding of the stimulus through infomax and its identification in the SORN subnetwork is a necessary condition to the completion of the BCI task in the IBEx model. In our simulations, since the stimulus lasts between 1 and 2 seconds and modulates the temporally filtered thalamic input, the timescale of the SORN firing rate adaptation must be neither too fast nor too slow to allow for the successful encoding of the stimulus without destroying the formation of synfire chains. The intrinsic plasticity parameter *η*_*IP*_ of the SORN model is tuned to allow for such constraint, since it affects the coupling of the unit firing rate to its target firing rate (Eq. 3).

Figure 5 shows the low-dimensional projection of SORN trajectories in the auditory cortex for values of *η*_*IP*_ spanning several orders of magnitude, so that the sensory stimulus can be adequately observed. When the intrinsic plasticity is too fast (i.e. for *η*_*IP*_ ≥ 10^−3^, the firing rate (FR) of each single unit of the SORN network is strictly forced to its target value of 10 Hz, and the stimulus cannot supply a sufficiently strong change in FR to be noticeable. In such scenario, the lack of variability in the FR prevents any form of rate-based coding. When the intrinsic plasticity is very slow (e.g. for *η*_*IP*_ *<* 10^−5^), the opposite effect is observed: the stimulus does not last long enough to create a noticeable change in the units FR. In such scenario, the coupling to the target FR is so loose that each unit is likely entering a slow cycle of continuous spiking and absolute silence which can take seconds or tens of seconds to complete, and refrain from encoding a short 1-2 seconds stimulus. An intermediate value of intrinsic plasticity of *η*_*IP*_ = 10^−4^ shows best encoding of the sensory stimulus and is used throughout the rest of the manuscript as other model components are integrated.

### iii. Information transmission across cortical areas

Having encoded the stimulus successfully in the sensory cortex, the next requirement concerned the reliable transmission of the stimulus to other cortices via synfire chains traversing the cortico-cortical feedback pathway to the motor cortex. This requirement implies that a sensory stimulus can prime arbitrary motor commands via synfire chains, which are then associated to a specific action, i.e. to control the neural activity of neurons in the pre-motor and motor cortices. In our model, SORN implements the cortico-cortical feedback pathway, and the transmission across brain regions amounts to the the propagation of synfire chains across SORN subnetworks. Once the intrinsic plasticity parameter *η*_*IP*_ is tuned to encode the stimulus (Section ii), the main parameter for propagating the stimulus to other SORN subnetworks is the spike-timing dependent plasticity rate *η*_*STDP*_. This parameter influences the creation of synfire chains since it dictates the temporal window at which pre-post and post-pre synaptic plasticity rules operate (see Eq. 2).

**Figure 5:**
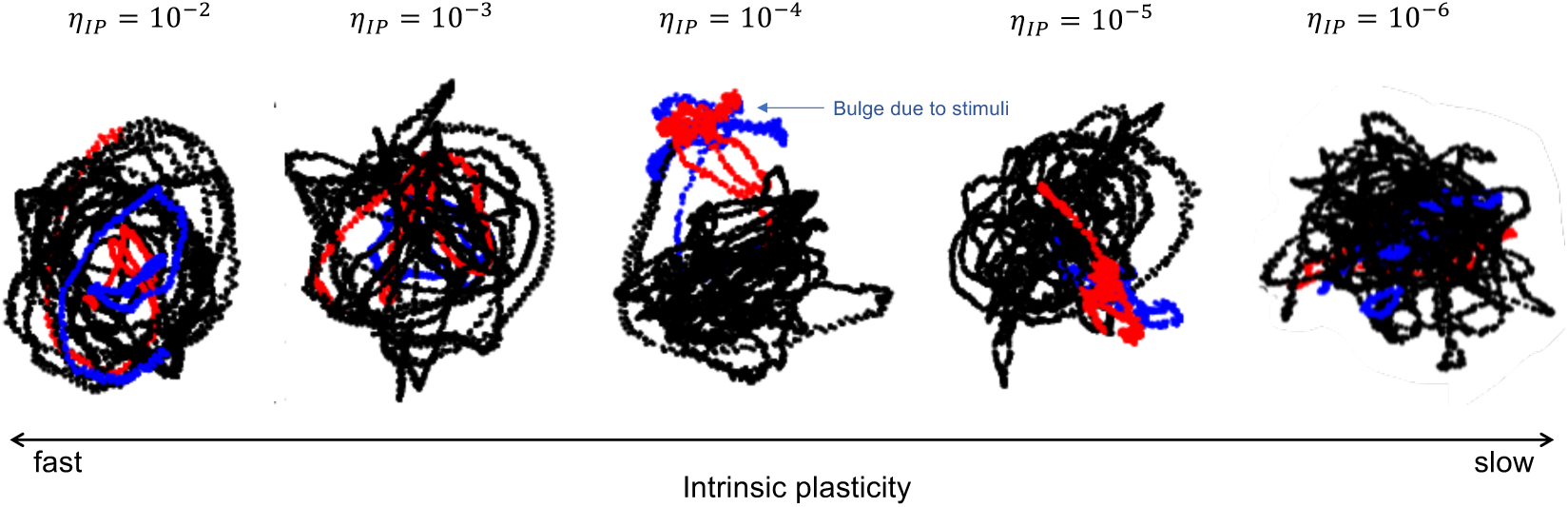
Adjusting L5 intrinsic plasticity η_IP_ for stimulus encoding in A1. Cortical trajectories of SVD projections for values of intrinsic plasticity η_IP_, also referred as homeostatic plasticity, spanning several orders of magnitude. Higher values of η_IP_ correspond to faster plasticity. Best encoding of stimulus is observed for η_IP_ = 10^−4^.

Figure 6 shows that, given *η*_*IP*_ = 10^−4^, for the original value of *η*_*STDP*_ = 4 · 10^−3^, the stimulus is successfully encoded by the sensory cortex but does not propagate reliably to connected regions (i.e. the bulge due to stimuli is not observable in adjacent SORN subnetworks). Slowing down of STDP by lowering *η*_*STDP*_ by an order of magnitude has a strong influence on the network structure, and can result in the loss of many connections, while strengthening the few that remain. This translates into more staggered cortical trajectories, but a more reliable transmission of the signal to neighbouring regions due to the strong connections remaining, as is observed in **Figure 6** for *η*_*STDP*_ = 4 · 10^−4^.

### iv. Striato-pallido-thalamic gating of cortical states

With an auditory stimulus that can be encoded in the sensory cortical region and propagated to nearby regions until reaching pre-motor and motor cortices, the third functional requirement of our analysis concerns the acquisition and selection of a motor command via the basal ganglia circuitry. The acquisition of such action requires reinforcement learning, which is part of the original model [9] but will not be addressed here. We focus instead on the downstream circuitry requirements of striato-pallido-thalamic projections to perpetuate an action decision (e.g. go / no-go) onto frontal (motor) cortices. Such decision (or selection) is supposedly initiated in the striatum and exerts its influence on the frontal cortices via the ventral thalamic nuclei under the modulation of basal ganglia inputs (the *“forward driving gate”* in the original paper [9]). The BCI learning task that we aim to reproduce *in silico* then requires a discernible cortical trajectory in motor areas under control of the auditory tone. It has been demonstrated that the acquisition of such motor commands requires cortico-striatal plasticity [8], implying a necessary role of the cortico-basal ganglia projections.

**Figure 6:**
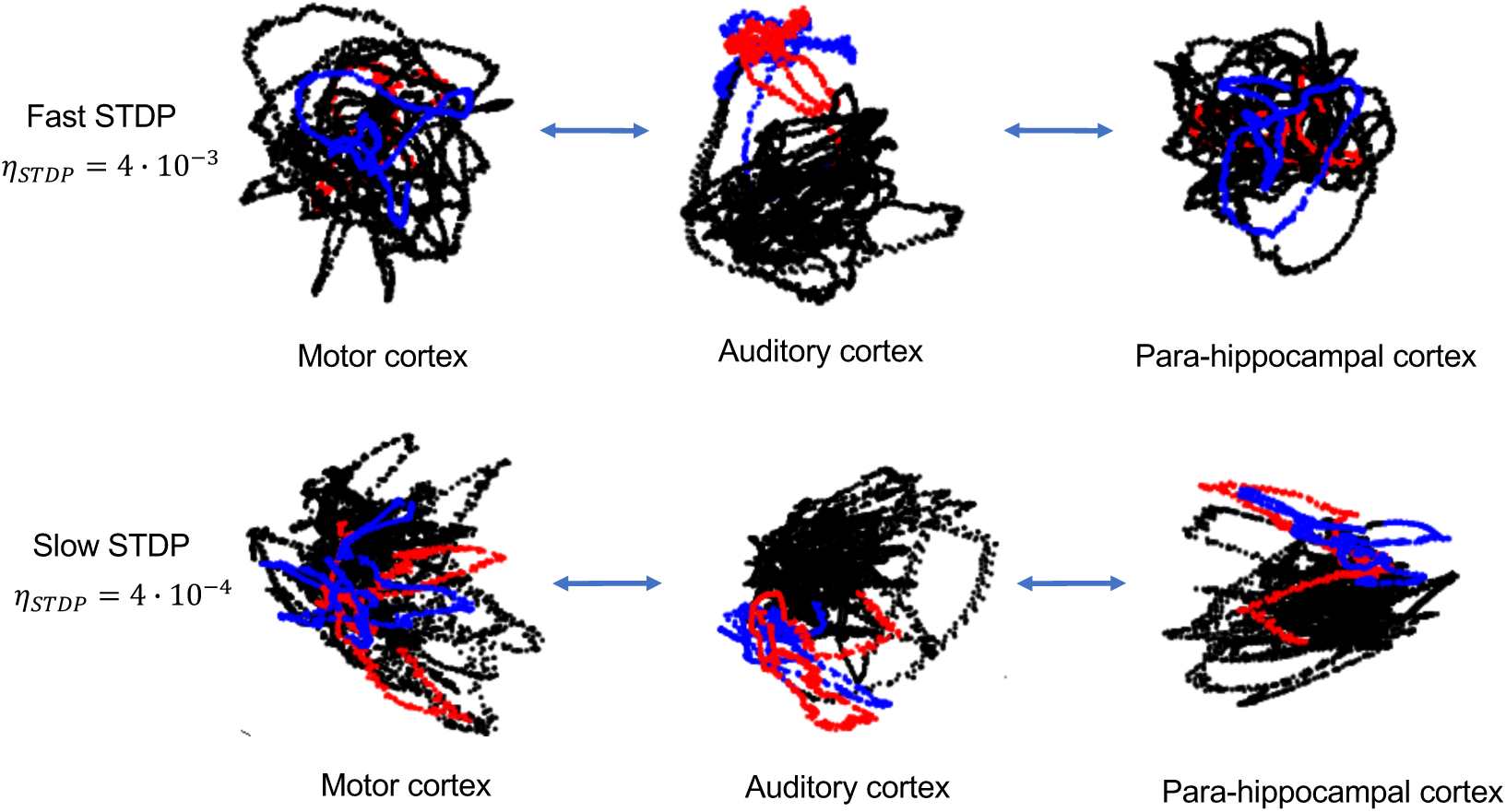
Adjusting L5 STDP for propagation of stimulus to other cortical areas. Cortical trajectories of SVD projections for values of spike-timing dependent plasticity (STDP) η_STDP_. Higher values of η_STDP_ correspond to faster plasticity. Best propagation of stimulus across areas is observed for η_STDP_ = 4 · 10^−4^.

The original IBEx model implements the striatum as a network for winnerless competition among inhibitory units, representing medium spiny neurons (the predominant neuronal type found in this structure). These units received spatially organized inputs from the cortical regions. In the winnerless inhibitory network [1], connections between neurons is random, with a connections probability set to 0.35, allowing recurrent bidirectional connections. Recent experimental studies on animal models observed that the topology of this inhibitory network is likely disrupted in disorders of the basal ganglia such as Huntington’s and Parkinson’s diseases, with a decrease of recurrent bidirectional connections [22]. Another more theoretical study stressed the need for some connectivity constraint in the striatum involving distance-based connectivity kernels [21].

We tested the consequence of such striatal network topology changes (removal of bidirectional connection, and distance-based connectivity constraint) onto the cortical dynamics of SORN via the striato-pallido-thalamic projections to frontal cortices. **Figure 7** shows an illustration of the different striatal topologies. In this experiment, we use a surrogate “open loop” striatal input in the form of a sequence of stimuli as a proxy for cortical dynamics instead of the “real” SORN input to calibrate the striato-pallido-thalamic system. Each striatal stimulus has a determined spatio-temporal structure or is drawn from a random distribution (**Figure 8**).

**Figure 7:**
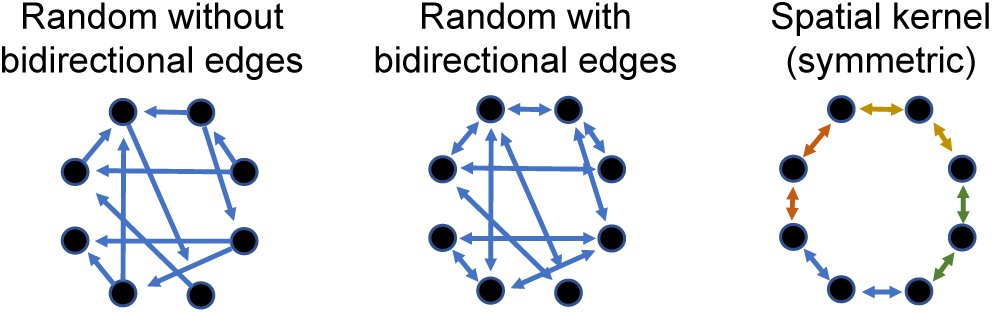
Striatal inhibitory network topologies. Striatal network connectivity can be random asymmetric (left), random symmetric (middle) or nearest neighbour using a symmetric spatial connectivity kernel (right). The connection probability of the random network is chosen to be 0.35, and the width of the connectivity kernel to span 10 units on each side in a circular network of 100 units.

**Figure 8:**
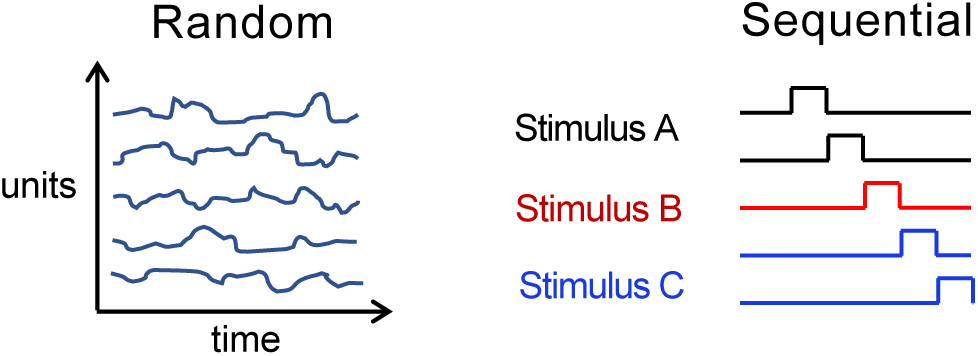
Surrogate striatal driving inputs for calibrating striato-pallido-thalamic gating. In the decoupled cortico-striatal system, surrogate striatal inputs are drawn from a random distribution (left) or follow a sequential activation (right), as a proxy to the SORN activity in the coupled system.

Figure 9 shows the motor cortex trajectories for the different scenarios of striatal connectivity and input patterns. The discernment of distinct cortical trajectories given the stimuli injected onto the striatum is best achieved using nearest neighbour striatal connectivity, whereby each striatal stimuli is confined to a region of the low-dimensional cortical space.

## IV. Discussion

We defined functional requirements necessary to perform a cognitive learning task involving the interaction of cortical and subcortical systems. We focused on cortical and thalamo-cortical dynamics that enabled stimulus encoding and propagation across brain regions, and in the striato-pallido-thalamic projections that permit to control frontal regions to modulate the stimulus. Important aspects of the system remain to be addressed to complete the task. Most notably, the BCI experiment is shown to rely on cortico-striatal plasticity [8], which is present in the original IBEx model under re-inforcement learning [9] but has been put aside while satisfying the preliminary requirements presented here. Modeling this learning task is particularly relevant to understand the pathophysiology of Huntington’s disease, which results in neurodegeneration among MSNs in the striatum, and the origin of which is believed to involve cortico-striatal dysregulation [19]. Future work in modeling cortico-striatal plasticity using realistic models of the basal ganglia will therefore take root among these existing models [5, 3].

**Figure 9:**
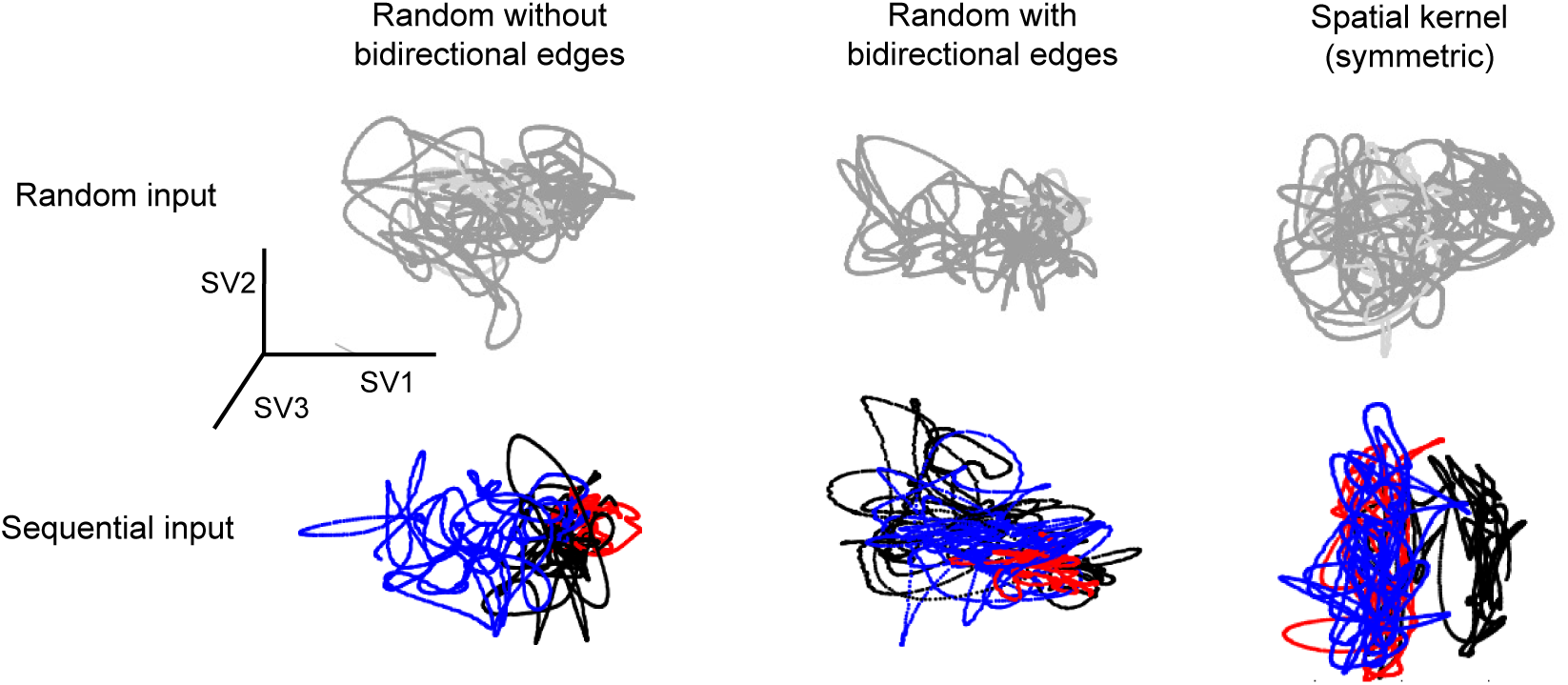
Effect of striatal dynamic on cortical trajectories. Low dimensional cortical trajectories of motor cortex M1 after SVD are shown for different striatal network structures and inputs. Striatal network structures are either random or nearest neighbour using a spatial connectivity kernel as depicted in **Figure 7**. For random networks, reciprocal connections between units can be enabled or disabled. Striatal driving input is either random (white noise) or a sequential activation pattern with distinct spatial organization as depicted in **Figure 8**.

It is notoriously difficult to control complex coupled non-linear systems like IBEx, with multiple learning elements (infomax), plasticity mechanisms (SORN), and network dynamics (winnerless) at play. Even setting aside reinforcement learning, probing the model within the functional requirements of this task was a challenge, and our attempt to reproduce a complex experimental finding using this complex system may seem vain. Still, useful insights were produced through the exercise.

### Future directions

Despite the complexity of the IBEx model, which we here attempted to reduce by analyzing decoupled sub-systems, some important components of the brain are left aside. For instance, our basal ganglia circuit does not incorporate the subthalamic nucleus (STN), which plays an important role in generating compulsive actions such as drug seeking behaviors, via the hyper-direct pathway through the basal ganglia, bypassing motor commands within the direct and indirect pathways. Also notable to the realization of the BCI learning task, the cerebellum may have a role to play in the refinement of motor commands, which may be useful to control the neuronal set recorded and ultimately in control of the auditory tone. Finally, the grand-loop hypothesis formulated by [9] spans the whole brain, but the model implemented here based on SORN is restricted by the number of feedforward clusters found in our particular SORN implementation, which in our simulation amounts to 4 cortical regions. A more elaborate cortical model using a brain atlas and its related connectivity matrix derived from white matter tractography would be welcome once the model is moved beyond proof-of-concept.

Our evaluation of cortical trajectories using SVD projection onto a low-dimensional space is not a sufficient metric for quantitative analysis of the framework, which remains to be developed. The eigenvalues of the SVD have been used to quantify the differences in cortico-thalamo-cortical dynamics with and without the supra-granular infomax activity, but are not sufficient to quantify specific differences among trajectories. We used visual inspection to give a qualitative appreciation of the cortical dynamics, and further quantitative measures of the span of these trajectories should be investigated, such as by measuring the Lyapunov exponent.

### Relation to other frameworks

Our cortico-thalamo-cortical system, modeled using SORN coupled to Infomax, contains features that can be mapped to Integrated Information Theory (IIT) [18]. IIT proposes a hierarchy of information measures linking past, current and future states of a network of mechanisms, to quantify the *amount of consciousness* in a system. In IBEx a SORN unit can be interpreted as a mechanism, and without intra-cortical infomax coupling, the cause-effect information *cei* can be inferred for each state that the system visits. Computing the minimum information partition *MIP* would remain a great challenge for non-trivial networks containing more than a few mechanisms (or units), as it requires dissecting the network hierarchically into multiple compositions of elementary mechanisms, over which time complexity scales exponentially. Interestingly, what is referred to in IIT as the *constellation of concepts* from which the integrated conceptual information Φ is derived, would correspond to the co-existing motor command in the current IBEx model with 4 cortical regions presented here. In this context, Φ may be seen as a proxy to the complexity of motor (and pre-motor) coordination.

Our framework is also compatible with the Global Neuronal Workspace hypothesis (GNW) [16], which is less mathematically oriented but more grounded in neuroanatomy than IIT. Most importantly, concepts developed in GNW directly applied to external consciousness (i.e. sensory awareness) are also transposable to internal consciousness (i.e. self-consciousness, introspection), and our attempt to model the BCI learning task falls at the interface between both. By formulating the functional requirements presented in this work and implementing them in the IBEx model, we contribute to the effort of understanding how complex, largely integrated cognitive functions emerge from neuronal circuits, and we provide a computational framework to explain them.

## V. Acknowledgments

This work was partly funded by the CHDI Foundation.

